# Next generation sequencing identifies a pattern of novel germline variants in early-onset colorectal cancer

**DOI:** 10.1101/2024.12.09.627474

**Authors:** Pierre Vande Perre, Ayman Al Saati, Bastien Cabarrou, Julien Plenecassagnes, Julia Gilhodes, Nils Monselet, Norbert Lignon, Thomas Filleron, Carine Villarzel, Laure Gourdain, Janick Selves, Mathilde Martinez, Edith Chipoulet, Gaëlle Collet, Ludovic Mallet, Delphine Bonnet, Rosine Guimbaud, Christine Toulas

**Affiliations:** Oncogenetics Laboratory, Oncopole Claudius Regaud, IUCT-Oncopole, Toulouse, France; DIAD team, INSERM U1037, Centre de Recherches en Cancérologie de Toulouse, Toulouse, France; Université de Toulouse, Université Toulouse III-Paul Sabatier, Toulouse, France; Biostatistics & Health Data Science Unit, Oncopole Claudius Regaud, IUCT-Oncopole, Toulouse, France; Bioinformatic Department, Oncopole Claudius Regaud, IUCT-Oncopole, Toulouse, France; Oncogenetics Department, Oncopole Claudius Regaud, IUCT-Oncopole, Toulouse, France; Pathology laboratory, Oncopole Claudius Regaud, IUCT-Oncopole, Toulouse, France; Oncology Department, Clinique Pasteur, Toulouse, France; CHU, Toulouse, France

**Keywords:** early-onset colorectal cancer, NGS panel, germline variants, molecular genetic of EOCRC

## Abstract

Early-onset colorectal cancer (EOCRC) incidence is increasing rapidly worldwide. However, the majority of EOCRCs are not substantiated by germline variants in the main colorectal cancer (CRC) predisposition genes (the “DIGE” panel). To investigate a potential genetic transmission of EOCRC (dominant, recessive and oligogenic hypotheses) and thus identify potentially novel EOCRC-specific predisposition genes, we conducted an analysis of 585 cancer pathway genes on an EOCRC patient cohort (n=87 patients diagnosed at ≤ 40 years of age, DIGE-) with or without a CRC family history. By comparing this germline variant spectrum to the GnomAD cancer-free database, we identified high impact variants (HVs) in 15 genes significantly over-represented in the EOCRC cohort. Among the 32 unrelated patients with a CRC family history (*i.e.* with a potentially dominant transmission pattern), nine presented HVs in ten of the genes tested, four of these genes had a DNA repair function. A potentially recessive transmission of EOCRC in patients without a CRC family history cannot be supported by our results nor can an oligogenic transmission.

We subsequently sequenced these 15 genes in a cohort of 82 late-onset CRCs (cancer diagnosis ≥50 years, DIGE-) and found variants in 11 of these genes to be specific to EOCRC. To evaluate whether variants in these 11 genes would allow to specifically detect EOCRC patients, we screened our patient database (n=6482), which only contained 2% of EOCRCs (DIGE-), and identified two other EOCRC cases diagnosed after the constitution of our cohort, with individual HVs in *RECQL4* and *NUTM1*. Altogether, we showed that 37.5% and 18.75% of heterozygous *NUTM1* and *RECQL4* HVs of our database were diagnosed with EOCRC.

Our work has identified a pattern of germline gene variants not previously associated with EOCRC. This paves the way to addressing the contribution of these variants to EOCRC risk and oncogenesis.

**Author Summary:** Early-onset colorectal cancer (diagnosed at ≤ 40 years of age) is a rare disease that can in part be explained by a hereditary genetic predisposition. To identify novel gene variants potentially associated with EOCRC risk, we analysed a panel of 585 genes in 87 patients with early-onset colorectal cancer unexplained by conventional genetic tests. This first analysis highlighted 15 genes of interest. To evaluate if this genetic profile is specific to early onset, we sequenced these 15 genes in a population of late-onset colorectal cancers (diagnosed after 50 years of age). Variants in 11 of these genes were specific to the early-onset population. To assess if this genetic pattern allows to identify other early-onset cases, we screened these genes in our whole database of 6482 patients and identified two new early-onset cases. Our results need to be confirmed, and validated in larger cohorts but pave the way for future research into early-onset colorectal cancer and the possibility of improving screening or treatment options for these patients and their family members.

## Introduction

Early-onset colorectal cancer (EOCRC) is a rare disease, but its incidence has increased by 2% every year over the past two decades (1, 2). A number of authors have therefore recommended generalised population-based colonoscopy screening from 45 years of age (3–5). EOCRC is characterised by late diagnosis/late-stage disease and left-sided primary tumours (1, 6–8). The increase in EOCRC incidence in high-income countries remains to be substantiated, as there are few epidemiological studies investigating this condition. EOCRC appears to be more prevalent in men and associated with a family history of colorectal cancer (CRC), inflammatory bowel disease, alcohol and tobacco use, as well as processed meat intake, a sedentary lifestyle and obesity (body mass index ≥30) (9, 10). Conversely, vegetable- and fruit-rich diets (11) and low-dose aspirin use (12, 13) appear to be protective.

It is generally accepted that 5-10% of familial CRCs are caused by a genetic predisposition transmitted by a Medelian pattern of inheritance (14). In line with the French national recommendations of the Genetics and Cancer Group (15), a consensus colorectal gene panel (DIGE panel) is therefore routinely used to identify germline pathogenic (PV) or likely pathogenic (LPV) CRC variants (in *APC, BMPR1A, CDH1, EPCAM, MLH1, MSH2, MSH6, MUTYH, PMS2, POLD1, POLE, PTEN, SMAD4* and *STK11*). Our team routinely screens EOCRCs against this DIGE panel and finds that between 70 and 80% of cases are not associated with germline PVs/LPVs in cancer predisposition genes. Several studies have used multi-gene testing (16–18) or whole exome sequencing (WES) (19–25) to identify the full spectrum of gene variants that specifically predispose to EOCRC (26). However, these studies have either tested small gene panels in large patient cohorts based on relatively non-selective inclusion criteria (e.g. 450 patients screened on a 25-gene panel (16), 315 patients on a 67- or 124-gene panel (17), 125 patients on a 94-gene panel (18)) or performed large molecular analyses (WES) on smaller cohorts (between 16 and 51 patients) (19, 20, 23, 25).

The aim of the current study was therefore to identify additional potentially novel germline EOCRC predisposition gene variants by sequencing 585 genes in a more selected EOCRC cohort that only includes cases testing negative on the DIGE panel genes (n=87). The EOCRC specificity of variant genes was examined by comparison to a late-onset, DIGE panel negative, CRC patient cohort (LOCRC, n=82), our local NGS patient database (n=6482) and the GnomAD NFE non-cancer cohort (n=51,377). A secondary aim based on EOCRC subcohorts with (n=32) and without (n=52) a CRC family history examined whether variants were inherited by a dominant, recessive and oligogenic transmission pattern.

## Results

### EOCRC cohort characteristics

The DIGE-, EOCRC (n=87) and LOCRC (n=82), cohorts had a median age at diagnosis of 34 and 62.5 years, respectively (Table 1). The EOCRC sex ratio (F/M) was 1.35 (p=0.197 compared to LOCRC cohort, see Table 1). The early-onset cohort characteristics were consistent with the literature in terms of distal disease (43.5% sigmoid or rectum involvement) and severity of disease at diagnosis (23.5% metastatic disease) (16–19, 21, 24, 25) and differed significantly from the LOCRC cohort (p <0.002). Most EOCRCs were microsatellite stable (MSS) or had low microsatellite instability (MSI-L) (76.2%) and were therefore mismatch repair proficient (pMMR), whereas LOCRCs had high microsatellite instability (MSI-H) and were mismatch repair deficient (dMMR).

**Table 1.** EOCRC (n=87) and LOCRC (n=82) patient characteristics. The number of patients (n), percentages (%, excluding any missing data) and medians (ranges) are listed for each category with p-values < 0.05 indicated in bold.

|  |  | **EOCRC**<br>**(n=87)** | **LOCRC**<br>**(n=82)** | **P-value** |
| --- | --- | --- | --- | --- |
| Median age at cancer diagnosis (years) |  | 34 | 62.5 | **<0.001** |
| (range) |  | (20 ;<br>40) | (50 ;<br>85) |  |
| Sex | Women | 50<br>(57.5%) | 39<br>(47.6%) | 0.197 |
|  | Men | 37<br>(42.5%) | 43<br>(52.4%) |  |
| Histology: adenocarcinoma | Liberkhunian | 46<br>(73.0%) |  |  |
|  | Colloid or mucinous | 10<br>(15.9%) |  |  |
|  | Signet ring cell carcinoma | 2<br>(3.2%) |  |  |
|  | Medullary | 0 |  |  |
|  | Mixed liberkhunian and<br>colloid | 3<br>(4.8%) |  |  |
|  | Mixed colloid and signet<br>ring cell | 1<br>(1.6%) |  |  |
|  | Mixed medullary and<br>signet ring cell | 1<br>(1.6%) |  |  |
|  | Missing data | 24 |  |  |
| Cancer by location | Small bowel | 4<br>(4.7%) | 2<br>(2.6%) |  |
|  | Appendix | 3<br>(3.5%) | 0 |  |
|  | Ascending colon | 18<br>(21.2%) | 39<br>(51.3%) |  |
|  | Transverse colon | 7<br>(8.2%) | 1<br>(1.3%) |  |
|  | Descending colon | 16<br>(18.8%) | 18<br>(23.7%) |  |
|  | Sigmoid | 19<br>(22.4%) | 6<br>(7.9%) |  |
|  | Rectum | 18<br>(21.2%) | 10<br>(13.2%) |  |
|  | Missing data | 2 | 6 |  |
| Location subclasses | "Proximal" (small bowel,<br>appendix, ascending<br>colon) | 25<br>(29.4%) | 41<br>(53.9%) | **0.002** |
|  | "Medial" (transverse and<br>descending colon) | 23<br>(27.1%) | 19<br>(25.0%) |  |
|  | "Distal" (sigmoid, rectum) | 37<br>(43.5%) | 16<br>(21.1%) |  |
| Colon without documented location |  | 2 | 6 |  |
| Metastatic CRC at primary diagnosis | Yes | 20<br>(23.5%) | 4<br>(5.5%) | **0.002** |
|  | No | 65<br>(76.5%) | 69<br>(94.5%) |  |
|  | Missing data | 2 | 9 |  |
| Tumour microsatellite instability | MSS or MSI-L | 48<br>(76.2%) | 13<br>(19.4%) | **<0.001** |
|  | MSI-H | 15<br>(23.8%) | 54<br>(80.6%) |  |
|  | Missing data | 24 | 15 |  |
| Immunohistochemistry<br>MMR protein expression | pMMR (MLH1, MSH2,<br>MSH6 and PMS2 normally<br>expressed) | 55<br>(79.7%) | 14<br>(19.7%) | **<0.001** |
|  | dMMR | 14<br>(20.3%) | 57<br>(80.3%) |  |
|  | Missing (or incomplete)<br>IHC | 18 | 11 |  |
| dMMR details | dMMR: MLH1 isolated<br>loss | 1 | 0 |  |
|  | dMMR: MSH2 isolated<br>loss | 0 | 0 |  |
|  | dMMR: MSH6 isolated<br>loss | 0 | 2 |  |
|  | dMMR: PMS2 isolated<br>loss | 0 | 10 |  |
|  | dMMR: MSH2/MSH6<br>combined loss | 5 | 10 |  |
|  | dMMR: MLH1/PMS2<br>combined loss | 8 | 31 |  |
|  | dMMR:<br>MLH1/MSH6/PMS2 loss | 0 | 1 |  |
|  | dMMR:<br>MLH1/MSH2/MSH6/PMS2<br>loss | 0 | 3 |  |
| Methylation of the MLH1 promoter | Yes | 2<br>(22.2%) | 20<br>(44.4%) | 0.283 |
|  | No | 7<br>(77.8%) | 25<br>(55.6%) |  |
|  | Missing data | 78 | 37 |  |
| BRAF V600E mutation | Yes | 3<br>(15.0%) | 11<br>(32.4%) | 0.160 |
|  | No | 17<br>(85.0%) | 23<br>(67.6%) |  |
|  | Missing data | 67 | 48 |  |
| CRC patients with a previous/subsequent<br>independent cancer history | Yes | 11<br>(12.6%) | 21<br>(25.6%) | **0.032** |
|  | No | 76<br>(87.4%) | 61<br>(74.4%) |  |
| Type(s) of independent cancer(s): | Colorectal | 2<br>(18.2%) | 3<br>(14.3%) |  |
|  | Small bowel NET | 1<br>(9.1%) | 0 |  |
|  | Cholangiocarcinoma | 1<br>(9.1%) | 0 |  |
|  | Prostate | 1<br>(9.1%) | 3<br>(14.3%) |  |
|  | Lung | 1<br>(9.1%) | 1<br>(4.8%) |  |
|  | Melanoma | 2<br>(18.2%) | 1<br>(4.8%) |  |
|  | Breast | 2<br>(18.2%) | 4<br>(19.0%) |  |
|  | Haematological malignancies (ALL, CLL) | 1<br>(9.1%) | 1<br>(4.8%) |  |
|  | Hodgkin's lymphoma | 1<br>(9.1%) | 0 |  |
|  | Skin lymphoma | 0 | 1<br>(4.8%) |  |
|  | Pineal dysgerminoma | 1<br>(9.1%) | 0 |  |
|  | Kidney | 0 | 3<br>(14.3%) |  |
|  | Pancreas | 0 | 1<br>(4.8%) |  |
|  | Bladder | 0 | 1<br>(4.8%) |  |
|  | Ovary | 0 | 3<br>(14.3%) |  |
|  | Adrenal cortex | 0 | 1<br>(4.8%) |  |
|  | Non-melanoma skin tumour | 0 | 2<br>(9.5%) |  |
|  | Endometrium | 0 | 1<br>(4.8%) |  |
|  | Thyroid | 0 | 2<br>(9.5%) |  |
| Family history of CRC or small bowel cancer | Yes | 32<br>(38.1%) | 33<br>(41.3%) | 0.680 |
|  | No | 52<br>(61.9%) | 47<br>(58.8%) |  |
|  | Missing data | 3 | 2 |  |
| If yes: | 1<sup>st</sup>-degree relative | 11<br>(34.4%) | 25<br>(75.8%) | **<0.001** |
|  | 2<sup>nd</sup>-degree relative | 24<br>(75.0%) | 13<br>(39.4%) | **0.004** |

EOCRC: early-onset colorectal cancer; LOCRC: late-onset colorectal cancer; CRC: colorectal cancer; MMR: mismatch repair; dMMR: deficient MMR (if expression of one of the four MMR proteins is lost); pMMR: proficient MMR (tumours expressing the four MMR proteins); MSS: microsatellite stable; MSI: microsatellite instability (*i.e.* MSI-H (high) if two or more markers are affected, MSI-L (low) if only one of the five markers is concerned); NA: not available; ALL: acute lymphocytic leukaemia; CLL: chronic lymphocytic leukaemia; NET: neuroendocrine tumour.

Most early-onset patients had no additional medical history of cancer, other than CRC (87.4%). Eleven patients had a previous or subsequent cancer history, independent of EOCRC. Patient EOCRC#19 (sigmoid adenocarcinoma at 31 years of age) presented with ascending colon adenocarcinoma at 52 and 56, dMMR, with loss of MLH1/PMS2, then a cholangiocarcinoma at 73 years of age. Patient EOCRC#26 (pMMR recto-sigmoid adenocarcinoma at 24) was subsequently diagnosed with a rectal adenocarcinoma at 26. Other independent tumour histories included small bowel neuroendocrine tumour (n=1), prostate cancer (n=1), lung cancer (n=1), melanoma (n=2), breast cancer (n=2), acute lymphoblastic leukaemia (n=1), Hodgkin lymphoma (n=1), pineal dysgerminoma (n=1) (EOCRC#1 had a melanoma and a lung cancer).

Only 38.1% of all EOCRCs had a CRC family history, 34.4% of these with an affected first-degree and 75.0% a second-degree relative. This distribution differed significantly in the LOCRC cohort, with 75.8% (p <0.001) and 39.4% (p=0.004) having an affected first- and second-degree relative, respectively. This was consistent with previous studies reporting a first-degree family history in 15.0% (3/20) and 13.6% (3/22) of EOCRC cases (19, 25).

### Shortlisting of germline high-impact variants in EOCRC patients

We sequenced 585 cancer predisposition genes/cancer pathway genes in 87 EOCRC patients and identified an average of 6425 variants per patient. To select variants that were preferentially detected in our EOCRC population we applied filters listed in Figure 1, we then compared both datasets from the EOCRC cohort and from the control GnomADv2 “non-cancer” subpopulation of non-Finnish European (NFE) origin (51,377 individuals). This approach identified 378 variants, including 10 truncating variants (TV, *i.e.* frameshift and non-sense variant), 4 missense variants identified as LPVs or PVs in the ClinVar database and 3 splice variants (SVs) in 15 different genes which were overrepresented (adjusted p-value <0.05) in our EOCRC population (Table 2 and Fig 2). After exclusion of 32 neutral (NVs) or likely neutral variants (LNVs), the remaining 329 variants were classified to be of unknown significance (VUS; Supporting information 3). These TVs/SVs/PVs/LPVs will hereafter be referred to as “high-impact variants” (HVs). Among the 15 EOCRC patients with HVs, none carried VUS in DIGE panel genes, with the exception of EOCRC#16 with a VUS in the exonuclease domain of *POLD1* (NM_002691.3:c.965G>A, p.(Arg322His), Supporting information 4).

**Fig 1.**
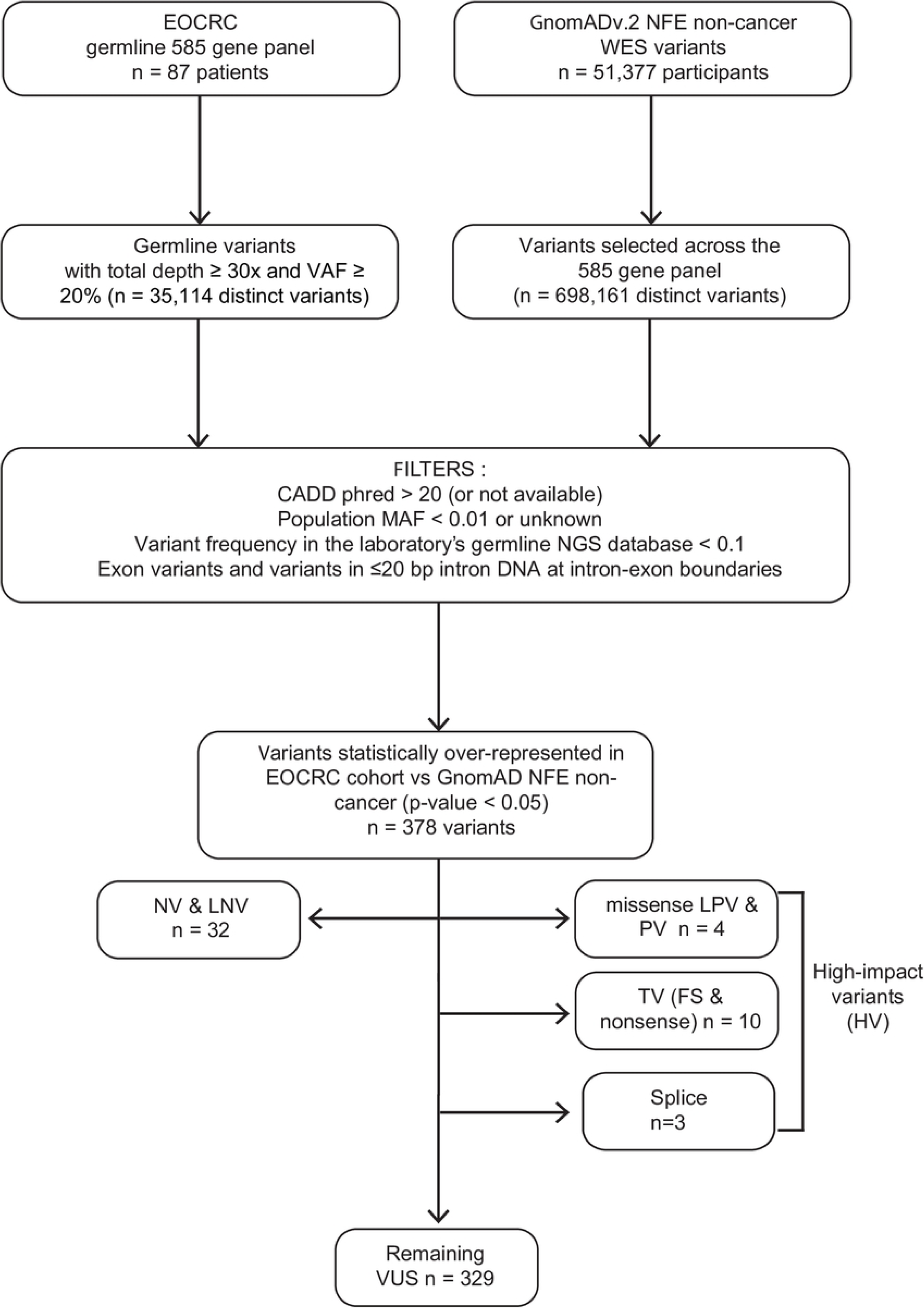
Strategy to identify EOCRC susceptibility genes for the monogenic dominant hypothesis. MAF: minor allele frequency; VAF: variant allele frequency; bp: base pair; NFE: Non-Finnish European; WES: Whole Exome Sequencing; EOCRC: early onset colorectal cancer; PV: pathogenic variant; LPV: likely pathogenic variant; VUS: variant of unknown significance; LNV: likely neutral variant; NV: neutral variant; TV: truncating variants (frameshift (FS) and nonsense variants).

**Fig 2.**
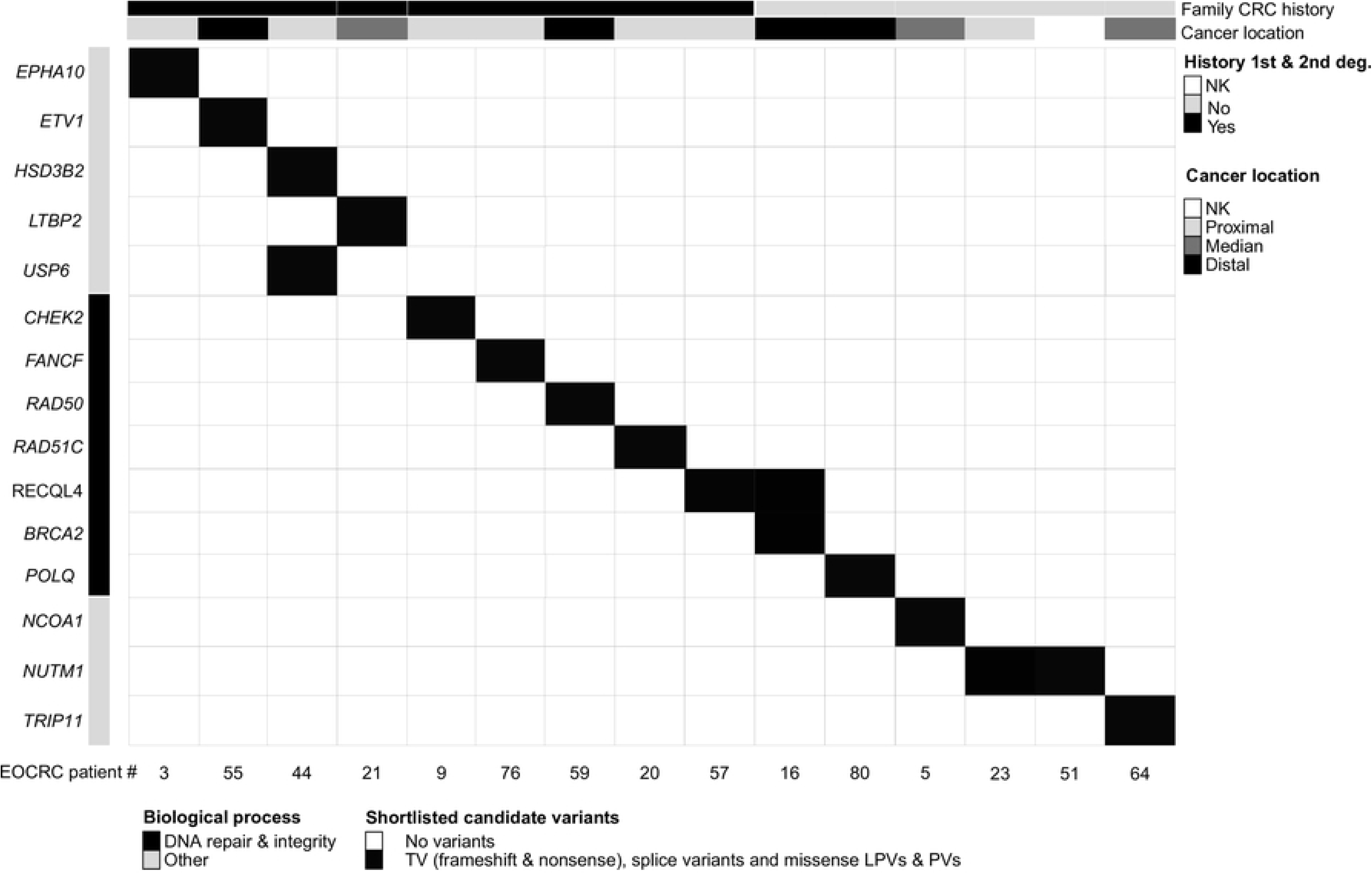
Clinical and molecular features of EOCRC patients with germline HVs. Each column corresponds to a patient (patient number specified). The upper section provides patient clinical features. Genes are grouped by biological processes (gene ontology). The upper section details patient clinical characteristics (1^st^ or 2^nd^ degree family history of colorectal cancer, anatomical digestive tract location of the disease). EOCRC: early-onset colorectal cancer; deg.: degree of kinship; NK: not known/not available; TV: truncating variants; PV: pathogenic variant; LPV: likely pathogenic variant.

**Table 2.**
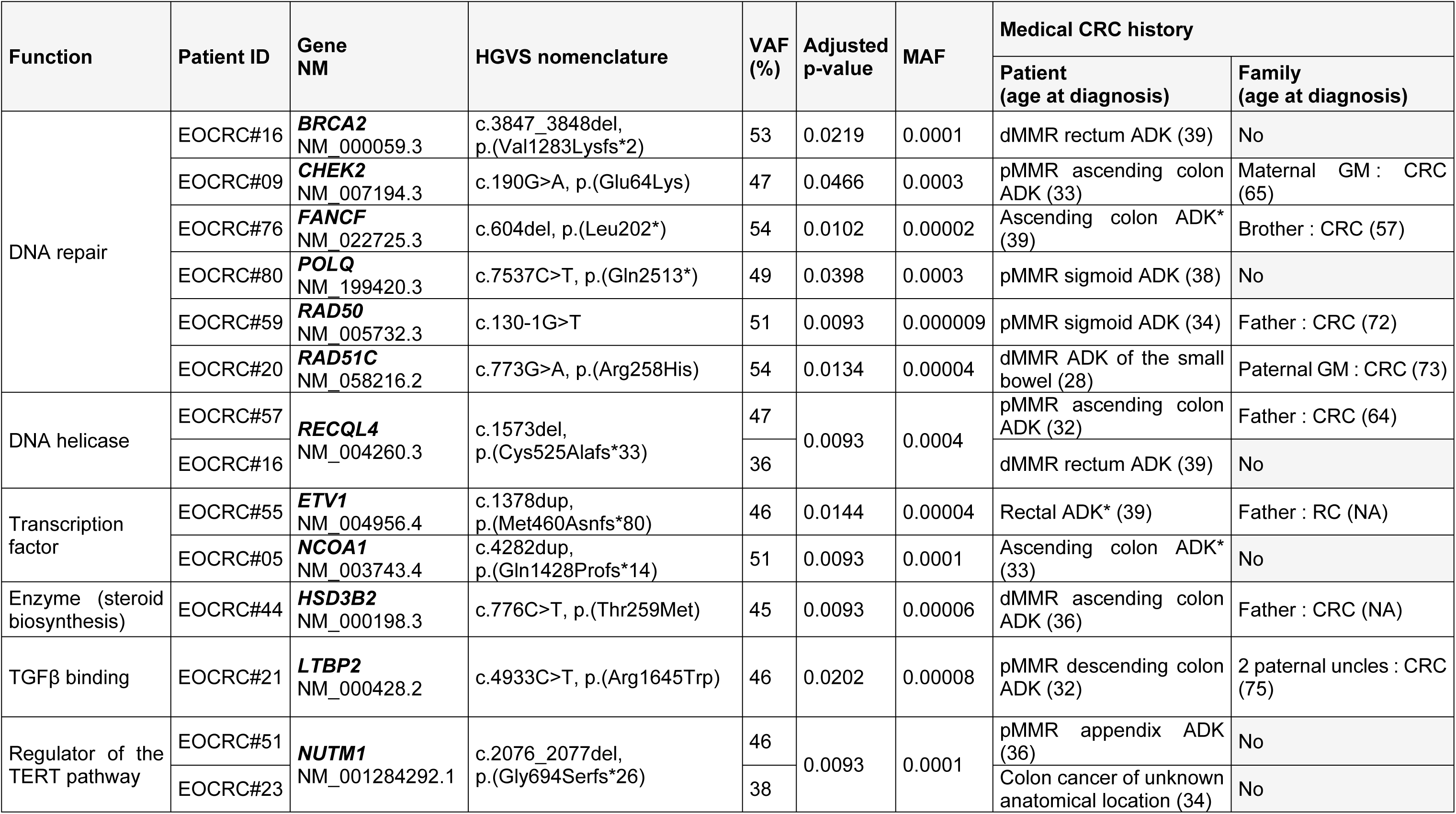
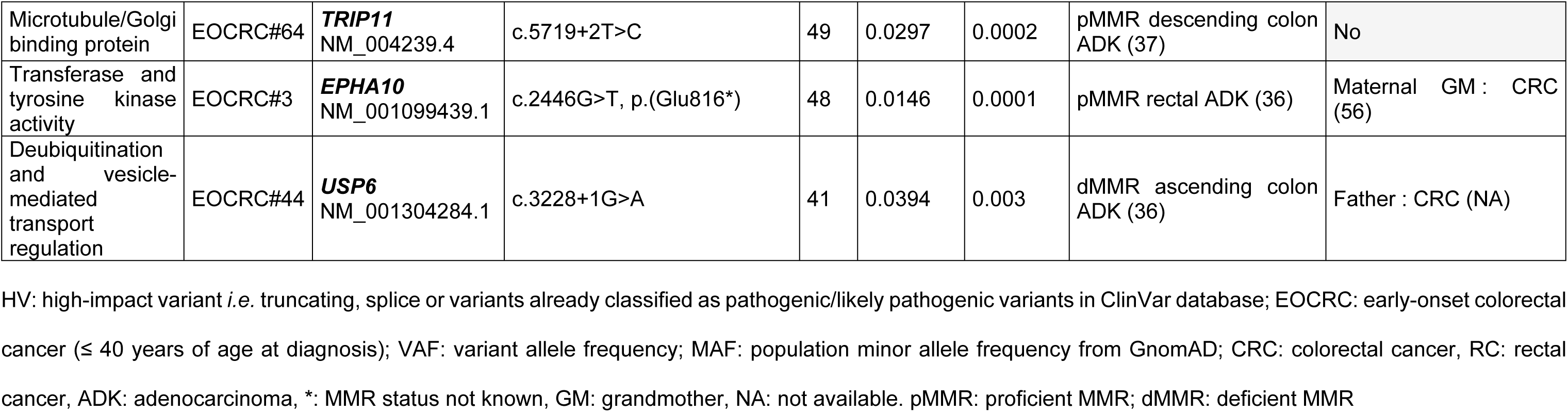
Molecular characteristics of germline HVs in EOCRC patients. P-values are specified for each variant.

| Function | Patient ID | Gene NM | HGVS nomenclature | VAF (%) | Adjusted p-value | MAF | Medical CRC history |  |
| --- | --- | --- | --- | --- | --- | --- | --- | --- |
|  |  |  |  |  |  |  | Patient (age at diagnosis) | Family (age at diagnosis) |
| DNA repair | EOCRC#16 | **BRCA2**<br>NM_000059.3 | c.3847_3848del,<br>p.(Val1283Lysfs*2) | 53 | 0.0219 | 0.0001 | dMMR rectum ADK (39) | No |
|  | EOCRC#09 | **CHEK2**<br>NM_007194.3 | c.190G>A, p.(Glu64Lys) | 47 | 0.0466 | 0.0003 | pMMR ascending colon ADK (33) | Maternal GM : CRC (65) |
|  | EOCRC#76 | **FANCF**<br>NM_022725.3 | c.604del, p.(Leu202*) | 54 | 0.0102 | 0.00002 | Ascending colon ADK* (39) | Brother : CRC (57) |
|  | EOCRC#80 | **POLQ**<br>NM_199420.3 | c.7537C>T, p.(Gln2513*) | 49 | 0.0398 | 0.0003 | pMMR sigmoid ADK (38) | No |
|  | EOCRC#59 | **RAD50**<br>NM_005732.3 | c.130-1G>T | 51 | 0.0093 | 0.000009 | pMMR sigmoid ADK (34) | Father : CRC (72) |
|  | EOCRC#20 | **RAD51C**<br>NM_058216.2 | c.773G>A, p.(Arg258His) | 54 | 0.0134 | 0.00004 | dMMR ADK of the small bowel (28) | Paternal GM : CRC (73) |
| DNA helicase | EOCRC#57 | **RECQL4**<br>NM_004260.3 | c.1573del,<br>p.(Cys525Alafs*33) | 47 | 0.0093 | 0.0004 | pMMR ascending colon ADK (32) | Father : CRC (64) |
|  | EOCRC#16 |  |  | 36 |  |  | dMMR rectum ADK (39) | No |
| Transcription factor | EOCRC#55 | **ETV1**<br>NM_004956.4 | c.1378dup,<br>p.(Met460Asnfs*80) | 46 | 0.0144 | 0.00004 | Rectal ADK* (39) | Father : RC (NA) |
|  | EOCRC#05 | **NCOA1**<br>NM_003743.4 | c.4282dup,<br>p.(Gln1428Profs*14) | 51 | 0.0093 | 0.0001 | Ascending colon ADK* (33) | No |
| Enzyme (steroid biosynthesis) | EOCRC#44 | **HSD3B2**<br>NM_000198.3 | c.776C>T, p.(Thr259Met) | 45 | 0.0093 | 0.00006 | dMMR ascending colon ADK (36) | Father : CRC (NA) |
| TGFβ binding | EOCRC#21 | **LTBP2**<br>NM_000428.2 | c.4933C>T, p.(Arg1645Trp) | 46 | 0.0202 | 0.00008 | pMMR descending colon ADK (32) | 2 paternal uncles : CRC (75) |
| Regulator of the TERT pathway | EOCRC#51 | **NUTM1**<br>NM_001284292.1 | c.2076_2077del,<br>p.(Gly694Serfs*26) | 46 | 0.0093 | 0.0001 | pMMR appendix ADK (36) | No |
|  | EOCRC#23 |  |  | 38 |  |  | Colon cancer of unknown anatomical location (34) | No |
| Microtubule/Golgi binding protein | EOCRC#64 | **TRIP11**<br>NM_004239.4 | c.5719+2T>C | 49 | 0.0297 | 0.0002 | pMMR descending colon ADK (37) | No |
| Transferase and tyrosine kinase activity | EOCRC#3 | **EPHA10**<br>NM_001099439.1 | c.2446G>T, p.(Glu816*) | 48 | 0.0146 | 0.0001 | pMMR rectal ADK (36) | Maternal GM : CRC (56) |
| Deubiquitination and vesicle-mediated transport regulation | EOCRC#44 | **USP6**<br>NM_001304284.1 | c.3228+1G>A | 41 | 0.0394 | 0.003 | dMMR ascending colon ADK (36) | Father : CRC (NA) |

HV: high-impact variant *i.e.* truncating, splice or variants already classified as pathogenic/likely pathogenic variants in ClinVar database; EOCRC: early-onset colorectal cancer ( $\leq 40$ years of age at diagnosis); VAF: variant allele frequency; MAF: population minor allele frequency from GnomAD; CRC: colorectal cancer, RC: rectal cancer, ADK: adenocarcinoma, \*: MMR status not known, GM: grandmother, NA: not available. pMMR: proficient MMR; dMMR: deficient MMR

Even though the relatively young age of the EOCRC cohort makes clonal haematopoiesis (CH) an unlikely rationale, we checked that the 15 EOCRC HVs were not included in the list of frequently mutated genes in CH (27) and that the variant allele frequencies (VAFs) were >35% (Fig 2 and Table 2). Patient EOCRC#09 carried a *CHEK2* gene variant included in the CH list, with a VAF of 47%, which seemed too high to be consistent with CH.

We pursued our analysis by successively investigating a dominant hypothesis for EOCRC, as well as recessive and oligogenic hypotheses.

### Investigating a dominant transmission pattern of EOCRC predisposition

We explored a monogenic dominant transmission pattern by statistically comparing variants of the EOCRC cohort (n=87) against those of the GnomAD NFE non-cancer population (n=51,377 individuals, see Fig 1). In our full EOCRC cohort, 15 patients carried statistically over-represented HVs (p<0.05) compared to the GnomAD NFE non-cancer control cohort.

Nine, of the 15 EOCRC patients with HVs, had a family history of CRC and carried HVs in 10 distinct genes (Fig 2 and Table 2), including genes involved in DNA repair pathways (*CHEK2*, *FANCF*, *RAD50* and *RAD51C*) and DNA integrity (*RECQL4*). Among these patients, EOCRC#9 (*CHEK2* variant) had a maternal family history of breast and pancreatic cancer (2^nd^ and 3^rd^ degree relatives). His maternal grandmother was diagnosed with both colorectal cancer and breast cancer at 65 years of age. *CHEK2* germline PVs/LPVs are considered as moderate-risk factors for breast cancer. To date no family *CHEK2* testing has been performed and the parental origin of the variant remains to be established. Patient EOCRC#57 (*RECQL4* variant) presented with a pMMR mucinous adenocarcinoma of the ascending colon at the age of 32 and her father was diagnosed with CRC at the age of 64. Although homozygous and compound heterozygous *RECQL4* pathogenic variants are known to cause recessive syndromes with overlapping features (Rothmund-Thomson syndrome (RTS, MIM #268400), RAPADILINO (MIM #266280) and Baller-Gerold syndrome (BGS, MIM #218600))(28), EOCRC#57 was heterozygous for the *RECQL4* PV/LPV (no evidence of compound heterozygosity) and did not exhibit any syndrome features. The remaining HVs in the EOCRCs with a family CRC history mapped to genes involved in steroid biosynthesis (*HSD3B2*), transcription regulation (*ETV1*) and other cellular pathways (*EPHA10*, *LTBP2*, *USP6*).

The remaining 6 EOCRC patients with HVs had no family history of CRC/small bowel cancer. They carried 7 variants in 6 distinct genes (Fig 2 and Table 2), including DNA repair (*BRCA2* and *POLQ*) and DNA integrity genes (*RECLQL4*). Patient EOCRC#16 with both BRCA2 (a PV inherited from her father) and *RECQL4* variants presented with a dMMR rectal adenocarcinoma at 39 years of age. The remaining variants mapped to genes involved in transcription regulation (*NCOA1*), Golgi apparatus trafficking (*TRIP11*) and possibly cell proliferation by modulation of TERT expression (*NUTM1*). In addition to the EOCRC initially reported at the age of 37, patient EOCRC#64 (*TRIP11* variant) was diagnosed with pineal dysgerminoma at the age of 21 and relapsed with a malignant germinoma. Patients EOCRC#23 (adenocarcinoma of the appendix at 36 years of age) and EOCRC#51 (colon cancer diagnosed at 38), from unrelated families, carried the same TV in *NUTM1*.

To investigate whether variants in these genes are associated with early onset, we screened these 15 genes in the later-onset CRC cohort (LOCRC, Table 3A and 3B). Four HVs (in the *CHEK2*, *FANCF*, *POLQ*, and *TRIP11 genes*), and 28 VUSs were identified in these 15 genes in LOCRC (also refer to Supporting information 5). Eleven genes were completely devoid of HVs in the LOCRC cohort. Six LOCR patients (7.3%) carried HVs in these genes, compared to 15 EOCRC patients (17.2 %). Patients LOCRC#69 and LOCRC#74 carried the same c.591del, p.(Val198Phefs*7) variant in *CHEK2*, but only LOCRC#69 had a CRC family history. Whilst 2 EOCRC patients carried the same HV in the *RECQL4* gene (one with and the other without a CRC family history) and 2 other EOCRC patients (both with a CRC family history) presented with the same HV in the *NUTM1* genes, none of the LOCRC patients had HVs in either of these genes.

**Table 3.**
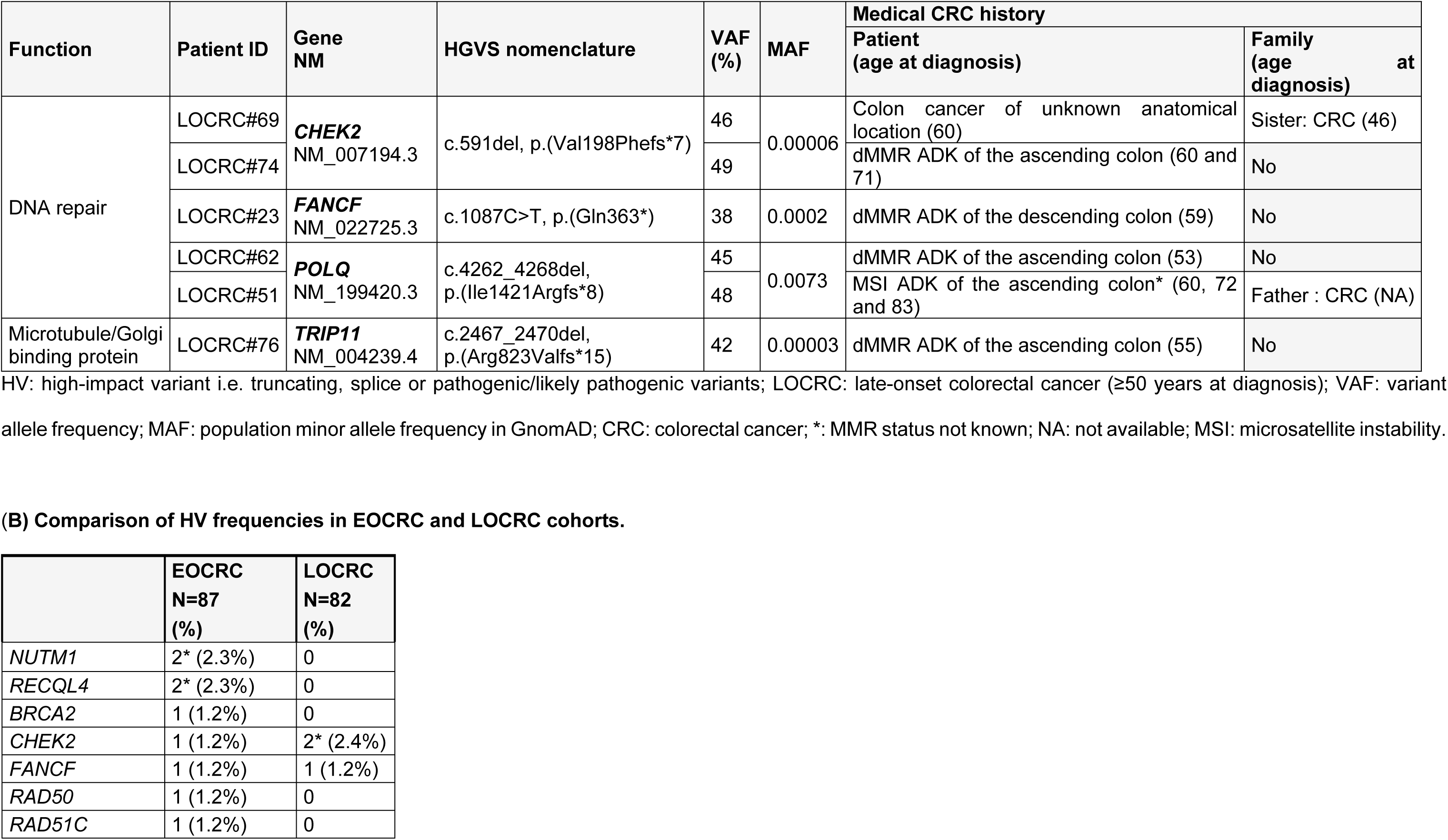
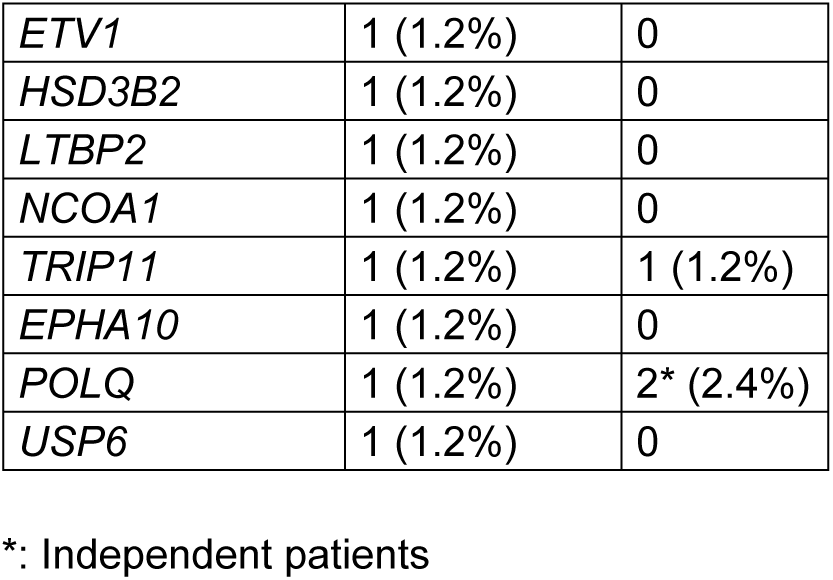
(A) Molecular characteristics of germline HVs in 15 genes of interest in the LOCRC cohort.

| Function | Patient ID | Gene NM | HGVS nomenclature | VAF (%) | MAF | Medical CRC history |  |
| --- | --- | --- | --- | --- | --- | --- | --- |
|  |  |  |  |  |  | Patient (age at diagnosis) | Family (age at diagnosis) |
| DNA repair | LOCRC#69 | **CHEK2**<br>NM_007194.3 | c.591del, p.(Val198Phefs*7) | 46 | 0.00006 | Colon cancer of unknown anatomical location (60) | Sister: CRC (46) |
|  | LOCRC#74 |  |  | 49 |  | dMMR ADK of the ascending colon (60 and 71) | No |
|  | LOCRC#23 | **FANCF**<br>NM_022725.3 | c.1087C>T, p.(Gln363*) | 38 | 0.0002 | dMMR ADK of the descending colon (59) | No |
|  | LOCRC#62 | **POLQ**<br>NM_199420.3 | c.4262_4268del, p.(Ile1421Argfs*8) | 45 | 0.0073 | dMMR ADK of the ascending colon (53) | No |
|  | LOCRC#51 |  |  | 48 |  | MSI ADK of the ascending colon* (60, 72 and 83) | Father : CRC (NA) |
| Microtubule/Golgi binding protein | LOCRC#76 | **TRIP11**<br>NM_004239.4 | c.2467_2470del, p.(Arg823Valfs*15) | 42 | 0.00003 | dMMR ADK of the ascending colon (55) | No |

HV: high-impact variant i.e. truncating, splice or pathogenic/likely pathogenic variants; LOCRC: late-onset colorectal cancer (≥50 years at diagnosis); VAF: variant allele frequency; MAF: population minor allele frequency in GnomAD; CRC: colorectal cancer; \*: MMR status not known; NA: not available; MSI: microsatellite instability.

**(B) Comparison of HV frequencies in EOCRC and LOCRC cohorts.**
|  | EOCRC<br>N=87<br>(%) | LOCRC<br>N=82<br>(%) |
| --- | --- | --- |
| *NUTM1* | 2* (2.3%) | 0 |
| *RECQL4* | 2* (2.3%) | 0 |
| *BRCA2* | 1 (1.2%) | 0 |
| *CHEK2* | 1 (1.2%) | 2* (2.4%) |
| *FANCF* | 1 (1.2%) | 1 (1.2%) |
| *RAD50* | 1 (1.2%) | 0 |
| *RAD51C* | 1 (1.2%) | 0 |
| *ETV1* | 1 (1.2%) | 0 |
| *HSD3B2* | 1 (1.2%) | 0 |
| *LTBP2* | 1 (1.2%) | 0 |
| *NCOA1* | 1 (1.2%) | 0 |
| *TRIP11* | 1 (1.2%) | 1 (1.2%) |
| *EPHA10* | 1 (1.2%) | 0 |
| *POLQ* | 1 (1.2%) | 2* (2.4%) |
| *USP6* | 1 (1.2%) | 0 |

\*: Independent patients

These analyses highlight differences between EOCRC and LOCRC patients, with HVs specifically identified in 11 genes of the early-onset CRC cohort.

To further explore whether variants in at least one of these 11 genes might be associated with EOCRC via a dominant transmission pattern, we examined our local germline NGS database for HVs in 10 of these genes (*BRCA2* was excluded) (Table 4). *BRCA2* PVs are not associated with colorectal cancer in the literature (29), and the frequency of *BRCA2* HVs in the EOCRC cohort does not appear to be higher than in the general population. *BRCA2* was therefore excluded from the subsequent analysis. The NGS database includes 6482 individuals tested for the germline predisposition to cancer hypothesis (mainly hereditary breast and ovarian cancer, CRC, pancreatic, prostate cancer and other cancers). HVs were identified in 103 “new” database patients (with a personal history of different types of cancers) and mapped to *NUTM1*, *RECQL4*, *RAD50*, *RAD51C*, *EPHA10*, *ETV1*, *LTBP2*, *NCOA1* and *USP6* genes (Table 4) but not to *HSD3B2*. No additional EOCRC case was identified among the screened database individuals with HVs in *RAD50, RAD51C, EPHA10, ETV1, LTBP2, NCOA1 and USP6*). In contrast, additional EOCRC cases were identified by screening the *NUTM1* and *RECQL4* genes. A more recent EOCRC patient (DIGE-) was identified with *RECQL4* c.2547_2548del, p.(Phe850Profs*33) TV, and is subsequently referred to as “EOCRC#88new”. She had dMMR rectal cancer at the age of 39 (MSH2/MSH6 loss) but no family history of CRC. The *NUTM1* TV c.3406C>T, p.(Arg1136*) was also identified in an additional EOCRC patient (DIGE-), thereafter referred to as “EOCRC#89new”. This patient presented with pMMR adenocarcinoma of the descending colon at the age of 31. Her mother and her only maternal aunt respectively developed CRC at 51 (pMMR) and 52 years of age. Finally, 37.5% (3/8) of heterozygous *NUTM1* TV carriers and 18.75% (3/16) of heterozygous *RECQL4* TV or SV carriers in the screened database, have been diagnosed with EOCRC.

**Table 4.** Results from a local NGS database screen (n= 6482 patients, of which 130 are EOCRCs without predisposing germline variants determined from routine testing against the DIGE panel) for HVs in 10 candidate genes (excluding *BRCA2*). Germline variants were filtered and classified as described in the “Patients and Methods” section. This screen identified 2 supernumerary EOCRCs (shown in bold): one with a heterozygous *NUTM1* non-sense variant and the other with a heterozygous *RECQL4* frameshift variant. These 2 “new” EOCRC patients (DIGE-) were included in the NGS database after the arbitrary 2021 cut-off date specified for the EOCRC cohort. LOCRC patients in this table were referred to the IUCT-O after 2021.

| Gene NM | HGVS nomenclature | Cancer personal history of carrier (number of carriers) |
| --- | --- | --- |
| NUTM1<br>NM_001284292.1 | c.303_304del, p.(Asp103Argfs*25) | EC (1, MSH6 mutation carrier) |
|  | c.2076_2077del, p.(Gly694Serfs*26) | EOCRC (2: #23 and #51); glioblastoma (1); OC (1) |
|  | c.2305del, p.(Glu769Argfs*3) | Cancer-free (1) |
|  | c.3106_3107del, p.(Lys1036Glyfs*7) | BC (1) |
|  | **c.3406C>T, p.(Arg1136\*)** | **EOCRC (1, supernumerary, "EOCRC#89new", diagnosed at 31 Y, DIGE-)** |
| RECQL4<br>NM_004260.3 | c.1048_1049del, p.(Arg350Glyfs*21) | BC (1) |
|  | c.1573del, p.(Cys525Alafs*33) | EOCRC (2: #57 and #16); polyposis (1); cancer-free patient (1); BC (3) |
|  | c.2269C>T, p.(Gln757*) | polyposis (1); BC (1); OC (1) |
|  | **c.2547_2548del, p.(Phe850Profs\*33)** | **EOCRC (1, supernumerary, "EOCRC#88new", diagnosed at 39 Y, DIGE-); MBC (1)** |
|  | c.2590C>T, p.(Gln864*) | Pancreatic cancer (1, PALB2 mutation carrier) |
|  | c.2755+1G>A | BC (1) |
|  | c.2994G>A, p.(Trp998*) | LOCRC (1, supernumerary, dMMR, diagnosed at 77 Y, DIGE-) |
| RAD50<br>NM_005732.3 | c.3229C>T, p.(Arg1077*) | BC (1) |
|  | c.541dup, p.(Ser181Phefs*7) | BC (1) |
|  | c.713_714insT, p.(Lys238Asnfs*8) | BC (1); OC (1); BC & OC (1); polyposis (1) |
|  | c.2165dup, p.(Glu723Glyfs*5) | BC (1) |
|  | c.3G>A, p.? | BC (1) |
|  | c.2938_2942del, p.(Leu980*) | NA (1) |
|  | c.1111dup, p.(Ile371Asnfs*14) | LOCRC (1, supernumerary, diagnosed at 66 Y, PMS2 mutation carrier) |
|  | c.354del, p.(Thr119Leufs*11) | BC (1) |
|  | c.3489_3495del, p.(Glu1164Glyfs*22) | BC (1) |
|  | c.2034del, p.(Gln678Hisfs*42) | BC (1); LOCRC (1, supernumerary, diagnosed at 68 Y, with polyposis, DIGE-); cancer-free patient (1) |
|  | c.130-1G>T | BC (1); EOCRC (1: #59) |
|  | c.2801dup, p.(Asn934Lysfs*10) | BC (1) |
|  | c.1281dup, p.(Gln428Thrfs*4) | BC (1) |
|  | c.1704del, p.(Tyr569Ilefs*29) | BC and Meningioma (1) |
|  | c.2985_2989del, p.(Glu995Aspfs*3) | BC (1) |
| RAD51C<br>NM_058216.2 | c.1026+5_1026+7del | OC (2) |
|  | c.31del, p.(Gln11Serfs*5) | BC (1) |
|  | c.709C>T, p.(Arg237*) | BC (1); OC (1) |
|  | c.965+5G>A | BC (2) |
|  | c.773G>A, p.(Arg258His) | EOCRC (1 : #20) |
|  | c.414G>C, p.(Leu138Phe) | BC (2); MBC (1); cancer-free patient (1); LOCRC (1, supernumerary, diagnosed at 50 Y, DIGE-) |
|  | c.577C>T, p.(Arg193*) | EC (1) |
|  | c.358dup, p.(Thr120Asnfs*35) | OC (1); cancer-free patient (1) |
|  | c.705+1G>A | OC (2) |
|  | c.732del, p.(Ile244Metfs*9) | OC (1); melanoma (1) |
|  | c.904+5G>T | BC (1) |
|  | c.656dup, p.(Leu219Phefs*33) | OC (1) |
|  | c.837+2T>C | BC (2) |
|  | c.837+1G>T | BC (1) |
|  | c.878del, p.(Asn293Ilefs*9) | Gastric cancer (1) |
| *ETV1*<br>NM_004956.4 | c.1378dup, p.(Met460Asnfs*80) | EOCRC (1: #55); MBC (1); pancreatic cancer (1) |
| *HSD3B2*<br>NM_000198.3 | c.776C>T, p.(Thr259Met) | EOCRC (1 : #44) |
| *LTBP2*<br>NM_000428.2 | c.4933C>T, p.(Arg1645Trp) | EOCRC (1 : #21) |
|  | c.468_469insG, p.(Thr157Aspfs*69) | BC (1) |
|  | c.4721-1G>C | BC (1) |
|  | c.709C>T, p.(Arg237*) | BC (1) |
|  | c.2789-1G>A | OC (1) |
| *USP6*<br>NM_001304284.1 | c.3228+1G>A | EOCRC (1 : #44); BC (2) |
|  | c.-1-2A>G | BC (2); OC (1); CRC (1, diagnosed at 47 Y, MSH2+) |
|  | c.3752dup, p.(Ser1252Glnfs*12) | BC (2); prostate cancer (1) |
|  | c.349C>T, p.(Gln117*) | BC (2); OC (1) |
|  | c.2828G>A, p.(Arg943His) | BC (1) |
|  | c.2828G>T, p.(Arg943Leu) | BC (1) |
|  | c.1337G>A, p.(Trp446*) | Cancer-free (1) |
|  | c.79C>T, p.(Arg27*) | BC (1) |
|  | c.4119C>G, p.(Tyr1373*) | BC (1); melanoma (1) |
|  | c.3028C>T, p.(Gln1010*) | CRC (1, diagnosed at 47Y, pMMR, DIGE-); BC (1) |
|  | c.2866C>T, p.(Arg956*) | BC (1) |
| *EPHA10*<br>NM_001099439.1 | c.2446G>T, p.(Glu816*) | EOCRC (1 : #03); BC (3); OC (1); cancer-free (1) |
|  | c.2920del, p.(Val974*) | OC (2) |
|  | c.1066C>T, p.(Arg356*) | BC (2) |
|  | c.851-1G>C | BC (1) |
| NCOA1<br>NM_003743.4 | c.4282dup,<br>p.(Gln1428Profs*14) | EOCRC (1 : #05) |
|  | c.3304-2A>G | BC (1) |

Y: years of age; NA: phenotype not available; HV: high-impact variant *i.e.* truncating, splice or pathogenic/likely pathogenic variants; BC: female breast cancer; MBC: male breast cancer; CRC: colorectal cancer; EOCRC: early-onset colorectal cancer ( $\leq 40$ Y); LOCRC: late-onset colorectal cancer ( $\geq 50$ Y); EC: endometrial cancer; OC: ovarian cancer.

### Investigating a monogenic recessive transmission pattern of EOCRC predisposition

To investigate the hypothesis of a monogenic recessive profile for EOCRC predisposition, we examined the subgroup of EOCRC patients without a CRC family history (“sporadic” cases, n=52) compared to variants in GnomAD control group as described in Fig 3. For the recessive homozygous hypothesis, variants (VUS, LPV or PV) were filtered to keep those with a high variant allelic frequency (VAF≥80%) in the analysis. For the recessive compound heterozygous hypothesis, we selected sporadic EOCRC patients carrying multiple variants (VUS, LPV or PV) (statistically over-represented in EOCRC vs GnomAD) in the same gene. However, we acknowledge that we do not have the necessary information to determine the different allele locations of the variants (*cis* or *trans*). Genes selected for the recessive hypothesis were also screened in the LOCRC cohort and in the full local NGS database with the same filtering process (Fig 3). Sequencing data from the 52 EOCRC patients with no family history were filtered and compared with GnomAD data as previously described (Fig 3). No significant homozygous variants (VAF≥80%) were identified in this sub-population of sporadic EOCRCs compared to the GnomAD NFE non-cancer. We then explored the hypothesis of potential compound heterozygosity in EOCRC patients without a CRC family history. Five EOCRC patients with no family history carried at least 2 variants of the same gene (Table 5): *CNTRL* (patient EOCRC#5 and EOCRC#72), *POLD1* (patient EOCRC#18), *RECQL4* (patient EOCRC#16) and *TCF12* (patient EOCRC#12) (Fig 3, Table 5). These 4 genes were then screened in the LOCRC cohort but none of the LOCRC patients were found to carry multiple variants in these genes.

**Fig 3.**
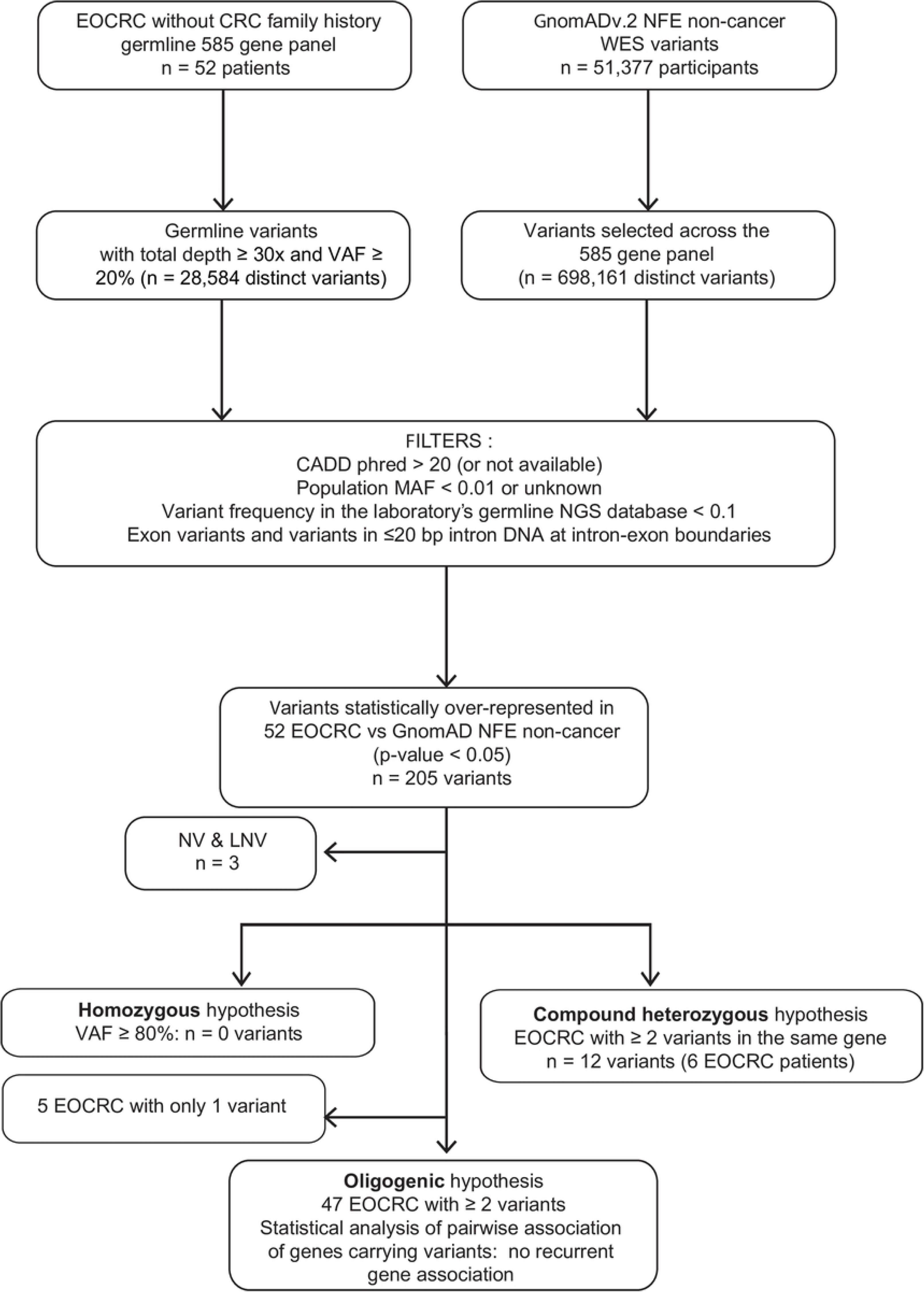
Strategy to identify EOCRC susceptibility genes for monogenic recessives (homozygous and compound heterozygous) and oligogenic approaches. MAF: minor allele frequency; VAF: variant allele frequency; bp: base pair; NFE: Non-Finnish European; WES: Whole Exome Sequencing; EOCRC: early onset colorectal cancer; PV: pathogenic variant; LPV: likely pathogenic variant; VUS: variant of unknown significance; LNV: likely neutral variant; NV: neutral variant; TV: truncating variants (frameshift (FS) and nonsense variants).

**Table 5.**
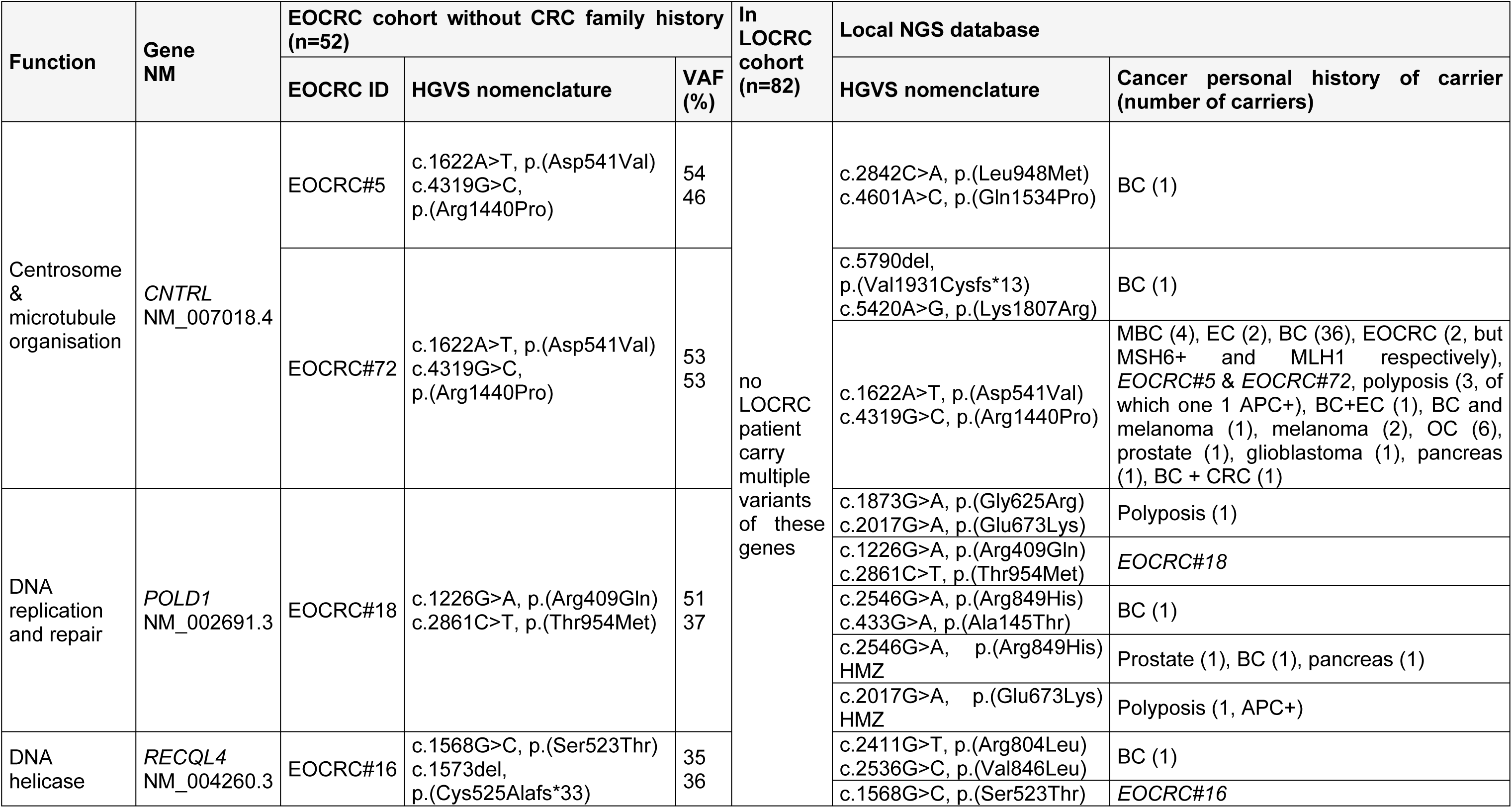
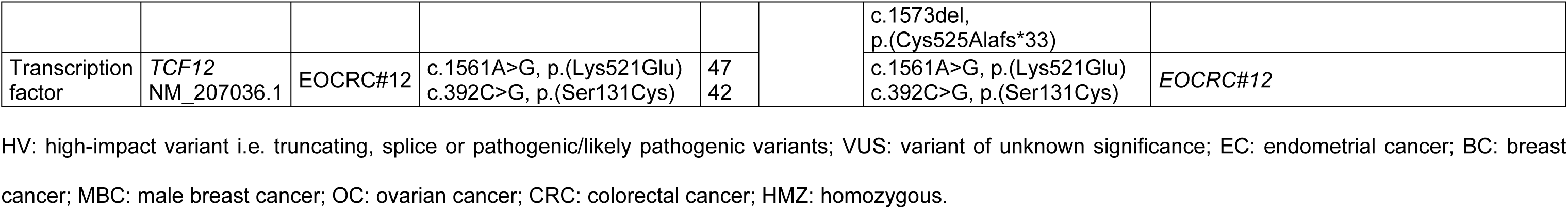
Recessive hypothesis: patients carrying multiple variants (HVs or VUSs) in the same gene. The LOCRCs and our local NGS database were screened for 6 genes of interest to identify patients carrying multiple variants in the same gene (after variant filtering as described in Fig 3).

We finally screened the whole NGS database for patients carrying multiple variants or homozygous variants in these genes. The same *CNTRL* variants found in EOCRC #5 and #72 were found in 61 patients diagnosed with a large spectrum of tumours, but excluding CRC (Table 5). Two patients were found to carry 2 variants in *POLD1* and four others in our NGS database were found to carry homozygous variant of *POLD1*, without CRC being over-represented in this population. Only one patient was identified with a double variant of *RECQL4,* with a breast cancer history and none of the features of a *RECQL4* recessive disorder, suggesting that these variants are located on the same allele. No other NGS database patient was found to carry multiple *TCF12* variants. Moreover, no patient in the NGS database was found to carry homozygous variants in *CNTRL*, *RECQL4* nor *TCF12*. These results strongly suggest that a recessive transmission of EOCRC risk can be reasonably dismissed in our population of 52 EOCRC patients without family history, at least for the 585 genes examined in this study.

### Investigating an oligogenic recessive transmission pattern of EOCRC predisposition

Finally, we wanted to investigate a possible recurrent pattern of gene association among sporadic patients (oligogenic hypothesis). We focused on variants (HVs and VUSs) that were significantly over-represented in sporadic EOCRCs vs GnomAD. Our genes of interest where those with variants detected in a least 3 EOCRC patients without a CRC family history (Fig 3). We next determined whether these genes occurred as pairs in the sporadic population (n=47, after ruling out 5 patients with a significant variant in only one gene).

Twelve variant-carrying genes were found in at least 3 patients each (Table 6). However, no recurrent association of variant-carrying-gene pairs was identified. The limited size of our population and the major limitations of large but targeted sequencing did not allow us to confirm or reject an oligogenic recessive transmission pattern of EOCRC predisposition in this study.

**Table 6.**
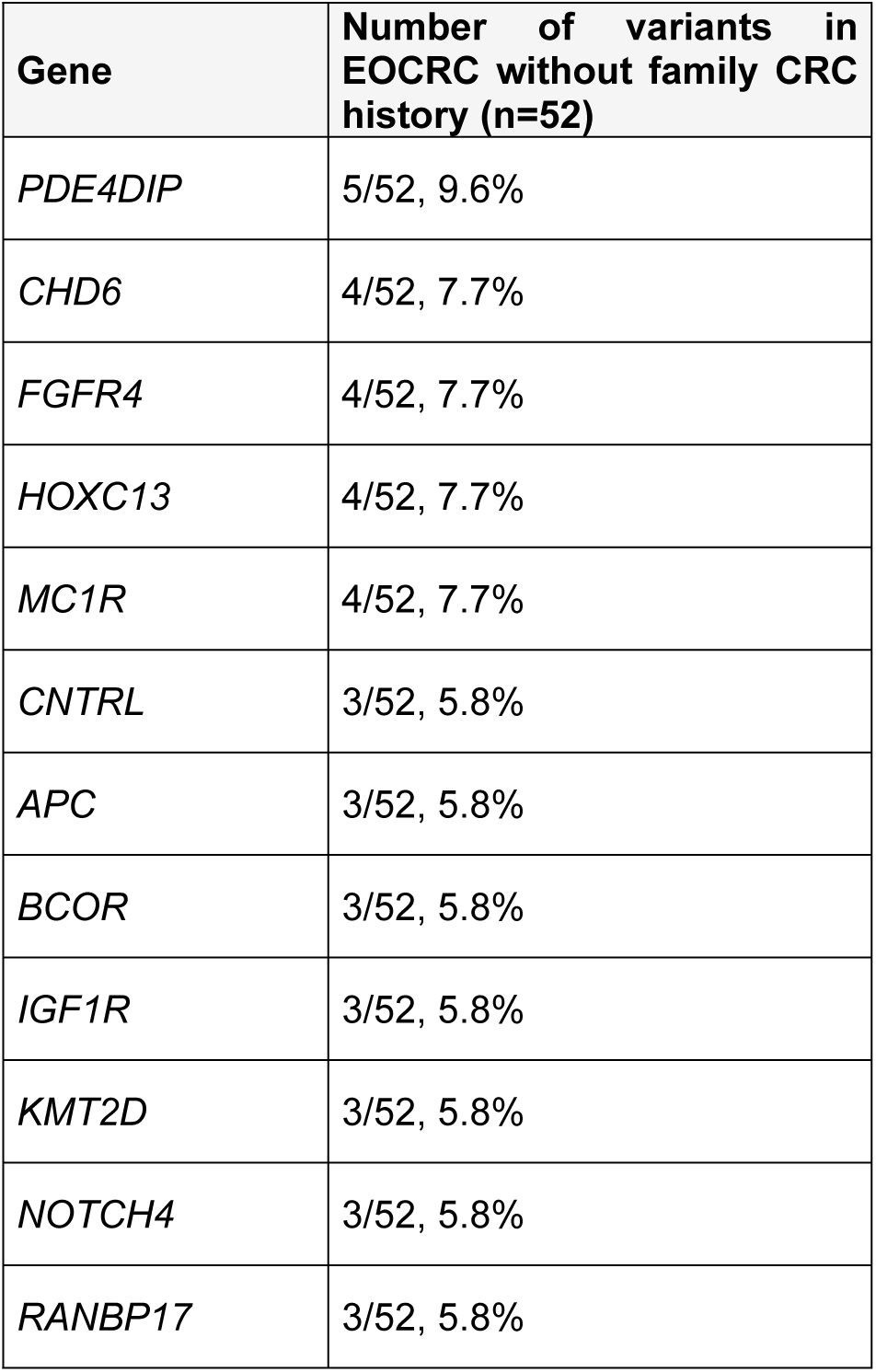
Oligogenic hypothesis: 12 variant-carrying genes, each found in at least 3 EOCRC patients without a CRC family history.

## Discussion

The aim of our study was to identify potentially novel gene variants that predispose to early onset colorectal cancer (EOCRC). We sequenced (NGS) a large panel of cancer and cancer pathway genes in a population of 87 EOCRC patients (with or without a CRC family history) diagnosed at ≤ 40 years of age and that tested negative for variants in the routine DIGE panel genes. We successively examined dominant, recessive and oligogenic transmission hypotheses as appropriate in the part of our cohort with (n=32) and without (n=52) a family history of CRC. We identified HVs that were significantly over-represented in our EOCRC cohort compared to the GnomAD non-cancer NFE population, in the following 15 genes: *BRCA2*, *POLQ*, *RAD50*, *RAD51C*, *RECQL4*, *CHEK2*, *FANCF*, *EPHA10*, *ETV1*, *HSD3B2*, *LTBP2*, *NCOA1*, *NUTM1, TRIP11* and *USP6*. Previous studies have investigated EOCRC populations and age cut-offs ranging from 40 to 55 years, either with small gene panels on larger populations (16–18) or by exome analysis on smaller populations (19–23, 25). These studies identified variants in known cancer susceptibility genes predominantly in DNA repair pathways (16–18, 22, 26) and in putative novel cancer genes such as *EIF2AK4* (20), *PTPN12* and *LRP6* (21). Our findings are consistent with these studies since 50% of our candidate (HV) genes are involved in DNA repair pathways (*BRCA2, CHEK2, FANCF, POLQ, RAD50, RAD51C* and *RECQL4*). Although *BRCA1* and *BRCA2* PVs/LPVs have previously been reported in 0.3 to 5% of EOCRC patients (16–18, 22, 24), the frequency of *BRCA* variants in EOCRC cohorts may simply reflect the frequency of these PVs/LPVs in the general population. A recent meta-analysis confirmed this conclusion by showing that carriers of *BRCA* mutations are not at increased risk of CRC (29). Our EOCRC#16 patient with the *BRCA2* PV also belongs to a high-risk breast cancer family. The evidence therefore suggests that PVs in *BRCA2* are not specific to EOCRC.

Even though our present study cannot formally confirm an association between HVs and EOCRC outcome, it is worth noting that most of the variants identified in DNA repair genes occurred in EOCRC patients with a 1^st^ or 2^nd^ degree family history of CRC (five out of seven patients or 71.4% of patients with DNA repair gene variants have a CRC family history compared to the 38% (n=32) of EOCRC patients that have a family history). Among the subgroup of 32 DIGE-EOCRC patients with a CRC family history, four additional patients carried variants in genes regulating TGF-β signalling (*LTBP2*), transcription (*ETV1*), cell-cell communication (*EPHA10*), steroid biosynthesis (*HSD3B2*) and vesicle-mediated transport (*USP6*) suggesting that cellular pathways other than DNA repair, are also involved in EOCRC risk.

In the 52 DIGE-, EOCRC patients without a CRC family history, the recessive monogenic transmission hypothesis was explored without identifying a homozygous variant of interest. Similarly, the search for compound heterozygosity was inconclusive and the gene association (oligogenic) hypothesis could not be confirmed. In this DIGE-EOCRC population without a family history of CRC, we also examined a dominant transmission hypothesis by identifying variants in EOCRC risk genes with incomplete penetrance. Among these 52 DIGE-EOCRC cases without familial involvement, six patients presented with HVs in six distinct genes, three patients with HVs in DNA repair genes (*BRCA2*, *POLQ* and *RECQL4*) and four patients with variants in genes regulating other cellular functions such as transcription (*NCOA1),* Golgi trafficking *(TRIP11)* and TERT regulation (*NUTM1*). *BRCA2* variants were excluded as explained earlier. *POLQ* is a DNA polymerase with helicase activity involved in DNA repair. *POLQ* germline variants have been reported in women with early-onset breast cancer (30) and are suspected to be linked to breast cancer but also to CRC (31) risk even though its role in CRC susceptibility has yet to be confirmed. As far as we know, *TRIP11* which maintains Golgi apparatus structure and interacts with thyroid hormone receptor beta has not been shown to be involved in CRC oncogenesis. In contrast, *NCOA1* also known as *SRC-1*/*RIP160*, is one of the main transcription co-activators of nuclear receptors whose dysregulation is involved in cancer initiation and progression (32). *NCOA1* also binds to β catenin a major actor of the Wnt pathway, extensively described in CRC oncogenesis (33).

We then attempted to determine whether this variant pattern is specific to the early onset of disease. The variant profile of these 15 genes of interest was analysed in a comparison cohort composed of 82 late-onset CRC patients (LOCRC, DIGE-). Eleven out of the 15 genes carried HVs exclusively in the DIGE-EOCRC cohort (no HVs in LOCRC cohort): *BRCA2*, *RAD50*, *RAD51C*, *RECQL4*, *EPHA10*, *ETV1*, *HSD3B2*, *LTBP2*, *NCOA1*, *NUTM1* and *USP6*. Even though our study is based on a very large but not exhaustive, panel of genes, these data underline the possibility of a distinct germline background in EOCRC and LOCRC.

We next assessed whether the analysis of these 10 genes of interest (*BRCA2* was excluded as explained above) could identify other EOCRC patients in a high-risk cancer NGS database. For this we screened our local NGS database that included data from 6482 patients, germline tested for various cancer predisposition hypotheses, including a total of 130 DIGE-, EOCRC patients, which included our study cohort of 87 EOCRC cases. HVs in *LTPB2*, *EPHA10*, *ETV1* and *NCOA1* were too rare to draw any useful conclusions. No other EOCRC patients were detected among individuals with HVs in *RAD50*, *RAD51C* and *USP6* genes (4.7% (1/21), 3.8% (1/26) and 4.5% (1/22) respectively). However, two other DIGE-EOCRC cases diagnosed after the recruiting period of our cohort carried truncated variants in *NUTM1* and *RECQL4*. *RECQL4* (RecQ Like Helicase 4) encodes a 3’ to 5’ DNA helicase that controls recombination, replication and DNA repair (34). Bi-allelic *RECQL4* defects are implicated in autosomal recessive inheritance syndromes (RTS, RAPALIDINO and BGS) but none of our *RECQL4* TVs heterozygous carriers showed clinical features of these conditions. Both *RECQL4* variants c.1573del, p.(Cys525Alafs*33) and c.2547_2548del, p.(Phe850Profs*33) (identified in the EOCRC#88new patient) are located in the helicase domain of the protein and are associated with recessive *RECQL4* disease (RTS and/or BGS) (28, 35–38). A retrospective study of self-reported family and medical history questionnaires of 123 heterozygous *RECQL4* carriers, from Rothmund-Thomson syndrome (RTS) families, found no significant difference in cancer risk compared to the general population but the location of the *RECQL4* variants were not well defined in this study (39).The *NUTM1* gene encodes the NUT protein (nuclear protein in testis, also called C15orf55), which is predominantly expressed in the testes but whose physiological function remains to be fully established. NUT may up-regulate telomerase reverse transcriptase (TERT) expression by binding GC boxes at the SP1 binding site of the *TERT* promoter (40). Tumours with high proliferation indexes, such as CRC, are known to have lost or reduced TERT expression (41). Patients EOCRC#51 and EOCRC#23 carry the same heterozygous *NUTM1* c.2076_2077del, p.(Gly694Serfs*26) variant which is predicted to shorten the C-terminus of the NUT protein by 440 amino-acids and removes the nuclear localisation sequence (NLS) (42). A supernumerary EOCRC#89new patient (DIGE-) was found to carry a heterozygous c.3406C>T, p.(Arg1136*) variant of *NUTM1* which is also predicted to truncate the last 36 amino-acids of the protein. Further studies are required to investigate the potential role of this C-terminal truncated NUTM1 protein in EOCRC oncogenesis. Interestingly, a previous genome-wide association study (GWAS) performed in a Korean population identified three loci associated with CRC, one of these included *NUTM1*, specifically the 20 amino acids before the C-terminus of NUTM1 (rs2279685, chr5:34649631) (43). Heterozygous TVs in *NUTM1* and *RECQL4* appear to be significantly over-represented, in our NGS database, among individuals with a history of EOCRC, with 38.5% (3/8) of *NUTM1* and 18.75% (3/16) of *RECQL4* TV carriers diagnosed with EOCRC. Considering that at the time this cross-sectional study was performed, the proportion of EOCRC patients (germline DIGE-) in our NGS database was 2% (130/6482), the EOCRC phenotype appears to be highly over-represented among the heterozygous *NUTM1* and *RECQL4* TV carriers.

To conclude, our study sequenced a panel of 585 genes which allowed to identify distinct germline variant patterns depending on whether the EOCRC patient had a CRC family history or not. These variant profiles also differed from those of a population with a later CRC diagnosis (LOCRC). In addition we independently identified other cases of DIGE panel negative EOCRCs from our whole NGS database that also contained variants in these genes. This work needs to be extended to determine the involvement of variants in genes of interest in altering the risk of EOCRC.

## Methods

### Cohorts

All CRC (or small bowel cancer) patients referred to the Oncogenetics Department (IUCT-O) between 2016 and 2021, with a negative germline test against the DIGE panel genes (DIGE-), were included in the analysis. We identified a DIGE-EOCRC cohort consisting of unrelated, adult patients (n=87) initially diagnosed with either early-onset colorectal or small bowel cancer (≤ 40 years of age at diagnosis), and a DIGE-cohort of late-onset CRC (or small bowel cancer) patients diagnosed after 50 years of age (LOCRC, n=82). All patients gave their written informed consent for the germline genetic analysis of cancer genes for both diagnostic and research purposes.

We retrospectively collated patient sex, age at diagnosis, cancer site, pathology, tumour phenotype when available (immunohistochemistry for mismatch repair (MMR) protein expression, microsatellite instability, BRAF V600E tumour mutation and *MLH1* promoter methylation status) as well as first-, second- and third-degree family history of cancer (maternal and paternal), from the medical records.

### MMR classification of tumour tissue

Tumour MMR and microsatellite status were determined from formalin-fixed colorectal or small bowel tumour tissue as previously described (44).

### Germline 585-gene panel sequencing

The Oncogenetics Laboratory of the IUCT-O (ISO15189 accredited) routinely sequences peripheral blood DNA with the germline panel [customised Comprehensive Cancer Panel (Roche, Switzerland)] of 585 cancer predisposition/cancer pathway genes (Supporting information 1). DNA was extracted from peripheral blood samples using QIAamp DNA Blood Mini kit (Qiagen, Germany), then captured using the KAPA library amplification kit (Roche) and sequenced on an Illumina Nextseq500 sequencer (Illumina, USA), as described elsewhere (45).

The GATK depth coverage tool was used to assess the mean sequencing depth for each gene of each individual sample (all exons and 20 bp of intron DNA at each intron/exon boundary). Target regions had a mean coverage of >98% at a ≥30x depth (Supporting information 2).

Data analysis for single nucleotide variants (SNVs) has been described elsewhere (45). Briefly, demultiplexing was performed with the bcl2fastq software (Illumina) and read alignment based on the GRCh37/hg19 genome assembly (using the *bwa mem* algorithm). Variants were identified using the HaplotypeCaller 3.3.0 and Varscan2 pipelines. Variants with a total sequencing depth of ≥30x and a variant allele frequency (VAF) ≥20% were selected for further analyses.

ANNOVAR and Ensembl Variant Effect Predictor were used to annotate the allele frequency of variants in the general population, *in silico* prediction scores (Combined Annotation Dependent Depletion – CADD-Phred score) (46), and other parameters, including the number of times each variant was detected in our local NGS oncogenetics database. Approximately 80% of the 6482 database patients were tested for breast/ovarian cancer, 10-15% for CRC or polyposis and 5-10% for other diseases (pancreas, prostate cancer, melanoma, etc…). The database also includes DNA from rare individuals who were cancer-free at the time of sampling but who are members of high cancer incidence families, with no surviving index patients.

### Variant analysis and classification

All coding (SNVs, small insertions and deletions) and intron variants (located in ≤20 bp intron DNA at intron/exon boundaries) were further evaluated. Rare variants were filtered by their predicted pathogenicity score (CADD-phred score >20 or not available), by their minor allele frequency (MAF) in GnomAD (whole base <1%) and were identified in fewer than 10% of NGS database samples (Fig 1 and 3). As copy number variants (CNVs) are not available in GnomADv2, these were not evaluated.

Variants that were significantly overrepresented in the DIGE-EOCRC group, compared to the control group, were classified according to the ACMG (American College of Medical Genetics and Genomics) recommendations (47). ACMG class 1 (NV) and class 2 (LNV) variants were excluded from the analysis. Statistically significant variants were confirmed by reviewing the BAM files with the Integrative Genomics Viewer (IGV). To be considered as a variant with a high predicted effect on splicing, the alteration had to be located in the exon or in the 2 base pairs immediately adjacent to either side of the intron/exon boundary (i.e., -2, -1 or +1, +2 relative to the splice junction) and the impact on splicing had to be predicted by both SpliceAI and SPIP pipelines (48, 49).

### Control population for statistical analysis of NGS data

The NGS data from the EOCRC cohort was compared to the germline data from the GnomADv2 “non-cancer” subpopulation of non-Finnish European (NFE) origin (51,377 patients) (50) and restricted to the 585-gene target regions (F/M ratio close to 0.5). It should be noted that approximately 85% of the control population was over 40 years of age.

### Statistical analysis

Categorical variables are expressed as frequencies and percentages and continuous variables as medians with ranges. Comparisons of the frequency of each variant between the DIGE-EOCRC and the GnomAD non-cancer NFE population were performed using the Fisher’s exact test with a Benjamini-Hochberg procedure for multiple testing. Comparisons between the DIGE-EOCRC and LOCRC groups were assessed using the Chi-squared or Fisher’s exact test for categorical variables and the Kruskal-Wallis test for continuous variables. All statistical tests were two-tailed and p-values <0.05 were considered statistically significant. Statistical analyses were performed using R (v4.1.2) and the STATA software (v18) (Stata Corporation, College Station, TX, USA). Gene ontology term scoring and pathway enrichment analysis, were performed with the g: Profiler.(51)

### Screening the local NGS database for candidate gene variants

The local NGS database (n=6482) was screened for the presence of candidate high impact variants (HVs) (i.e., TVs, SVs and variants known to be PVs/LPVs) identified in the EOCRC cohort. Variants were filtered and classified using the criteria described above.

## Acknowledgments

We would like to thank Mrs Lydia Loncle, Mrs Pauline Dejean, Mrs Aurelia Da Cruz, Mrs Isabelle Gitlaw and all the technicians from the Toulouse Oncogenetics Laboratory, without whom this work would not have been possible. We would also like to thank the French association Comminges Sans Frontières for funding a part of the ancillary expenses associated with this work.

## Supporting information captions

**SI Table 1: List of the 585 germline panel genes.**

**SI Table 2: Germline 585 gene panel sequencing coverage and quality statistics of EOCRC samples.** As NGS has been performed for germline medical purposes, the minimum threshold of coverage is ≥30x. Read alignment was made to the GRCh37/hg19 genome assembly. * targeted bases = the 585 gene panel (gene list described in supporting information 1)

**SI Table 3: All 329 VUSs identified in the EOCRC cohort compared to the GnomAD NFE non-cancer population with adjusted p-value <0.05.** VAF: variant allele frequency

**SI Table 4: VUSs identified in the 15 EOCRC patients with germline high-impact variants.** VAF: variant allele frequency

**SI Table 5: VUSs identified in the LOCRC cohort among the 15 genes found to carry high-impact variants in EOCRC.** VAF: variant allele frequency

## Financial Disclosure Statement

The authors received no specific funding for this work.

## Ethical approval

As this study is a completely retrospective study with no implications for human participants, no ethics approval was required. All data analyses (germline, tumour and patient characteristics) were generated for healthcare purpose and analysed retrospectively. This study complies with declaration n°2206723 of the University Hospital of Toulouse to the Commission Nationale de l’Informatique et des Libertés (CNIL).

## Competing Interests

The authors declare no competing interests in relation to this work.

## Notes

### Competing Interest Statement

The authors have declared no competing interest.

